# Sensitivity of radio-photoluminescence glass dosimeters to accumulated doses

**DOI:** 10.1101/2020.06.04.133900

**Authors:** Dong Wook Kim, Jiwon Sung, Jaeman Son, Han-Back Shin, Min-Joo Kim, Yu-Yun Noh, Hojae Kim, Min Cheol Han, Jihun Kim, Su Chul Han, Kyung Hwan Chang, Hojin Kim, Kwangwoo Park, Myonggeun Yoon, Jinsung Kim, Dongoh Shin

## Abstract

**Background:** This study investigated the effect of accumulated doses on radio-photoluminescence glass dosimeters (RPLGDs) from measurements involving mega-voltage photons.

**Methods:** Forty-five commercially available RPLGDs were irradiated to estimate their dose responses. Photon beams of 6, 10, and 15 MV were irradiated onto the RPLGDs inside a phantom, which were divided into five groups with different doses and energies. Groups 1 and 2 were irradiated at 1, 5, 10, 50, and 100 Gy in a sequential manner; Group 3 was irradiated 10 times with a dose of 10 Gy; and Groups 4 and 5 followed the same method as that of Group 3, but with doses of 50 Gy and 100 Gy, respectively.

**Results:** For the annealed Group 1, RPLGD exhibited a linearity response with variance within 5%. For the non-annealed Group 2, readings demonstrated hyperlinearity at 6 MV and 10 MV, and linearity at 15 MV. Following the 100 Gy irradiation, the readings for Group 2 were 118.7 ± 1.9%, 112.2 ± 2.7%, and 101.5 ± 2.3% at 6, 10, and 15 MV, respectively. For Groups 3, 4, and 5, the responsiveness of the RPLGDs gradually decreased as the number of repeated irradiations increased. The percentage readings for the 10th beam irradiation with respect to the readings for the primary beam irradiation were 84.6 ± 1.9%, 87.5 ± 2.4%, and 93.0 ± 3.0% at 6 MV, 10 MV, and 15 MV, respectively.

**Conclusions:** The non-annealed RPLGD response to dose was hyperlinear for the 6 MV and 10 MV photon beams but not for the 15 MV photon beam. Additionally, the annealed RPLGD exhibited a fading phenomenon when the measurement was repeated several times and demonstrated a relatively large fading effect at low energies than at high energies.

## Background

Radiation therapy, along with surgery and chemotherapy, plays a critical role as a main treatment approach for cancer patients (1, 2). Furthermore, radiation therapy trends are shifting from traditional three-dimensional conformal radiation therapy (3DCRT) techniques to complex radiation delivery techniques, such as intensity-modulated radiation therapy (IMRT), volumetric modulated arc therapy (VMAT), and helical tomotherapy (TOMO), all of which deliver more precise and localized dose distributions (3, 4). Because radiation therapy techniques have become more complex and sophisticated, the importance of proper quality control (QC) and quality assurance (QA) for precise patient care has increased (5-7), thereby increasing the importance of accurate in vivo and phantom dose measurements.

In vivo measurement is an important procedure for verifying that, during treatment, a radiation dose has irradiated the patient accurately according to the treatment plan. In vivo measurements are performed at several locations, with these measurements requiring high spatial resolution, high sensitivity, and low dose influence, over a broad range of doses. Silicon diodes, Gafchromic films, metal–oxide–semiconductor field-effect transistors (MOSFETs), and thermoluminescence detectors (TLDs) are all currently used as in vivo dosimetric tools (8-14). These dosimetric sensors each have their own inherent advantages and disadvantages. TLDs have advantages, such as small detector size, good reproducibility, and radiation sensitivity, but also have disadvantages in terms of angular dependence, energy dependence, and relatively large workloads in calibration and measurement procedures (15). In comparison with TLDs, diodes and MOSFETs, have an advantage in terms of eliciting immediate responses, but these detectors have relatively high initial costs, and exhibit fading effects after dose limitations (8). Meanwhile, external beam therapy 3 (EBT3) film, the representative Gafchromic film that is currently in use, has an advantage in terms of being thin, and also permits two-dimensional dose distribution; however, it has a relatively long saturation time, and exhibits high uncertainty at low doses (10). In 1999, the optically stimulated luminescence dosimeter (OSLD) was introduced by McKeever et al.; it comprises crystalline aluminum oxide doped with carbon (Al_2_O_3_:C), and is characterized by high radiation sensitivity, good dose linearity, and a low effective atomic number, and does not suffer from fading (16). Although they suffer signal losses of 1–2%, OSLDs have an advantage in terms of allowing repeat readings, accumulated readings, sensor identification QR codes, having relatively short reading procedures (∼10 min), and having simple read outs that use light instead of heat, which reduces the risk of damage to the detectors. With these advantages, the OSLD is rapidly becoming important in the field of in vivo dosimetry (17).

A commercially available radio-photoluminescence glass dosimeter (RPLGD, GD-302M, Asahi Techno Glass Co., Shizuoka, Japan) has been introduced in Japan as an alternative to TLDs. Similar to OSLDs, RPLGDs are advantageous in that they allow repeat readings, accumulated readings, sensor identification numbers, have relatively short reading procedures, and use the same simple read outs as those of OSLDs (18, 19). In 2009, Lee et al. presented the dosimetric characteristics and performance of the RPLGD for environmental exposure situations (20). They found that the RPLGD fading was approximately 1% within 30 days of being exposed to approximately 0.6, 6, and 20 mGy. Meanwhile, in a later study in 2019, Shehzadi et al. discovered, via a repeatability test, that the obtainable statistical uncertainty of the RPLGD was within 1% (21). They estimated the amount of deviation in the RPLGD reading after irradiation with approximately 1, 3, 5, and 9 Gy using a therapy-level Co-60 gamma ray beam. The RPLGD has relatively good reproducibility and exhibits low energy dependence at energies higher than 200 keV. Additionally, the rod-shaped RPLGD, which has a round cross section, may have a relatively small incident-beam angular dependence on the orthogonal direction of the sensor. In comparison with TLDs or OSLDs, the RPLGD has the advantage of being easy and safe to use inside the human body, because of its low toxicity and thin rod shape.

In recent years, RPLGDs have become important in the field of in vivo dosimetry. However, no study on the accumulative dose response of RPLGDs has been conducted thus far. Therefore, in this study, we evaluated and reported on the accumulated dose response of RPLGDs for mega-voltage therapeutic X-ray beams.

## Methods

### RPLGD

The commercially available RPLGD used in this study (GD-302M, Asahi Techno Glass Co., Shizuoka, JAPAN) was a colorless and transparent glass rod with a diameter of 0.15 cm and a length of 0.85 cm. The by-weight composition of the RPLGD was 51.16% of O, 31.55% of P, 11.00% of Na, 6.12% of Al, and 0.17% of Ag (22). The density and effective atomic number of RPLGD were 2.61 g/cm^3^ and 12.039, respectively (22). The RPLGD used had a thin plastic cover that was 0.28 cm in diameter and 0.13 cm in length. A dose estimate for the RPLGD in the reader (FGD-1000; Asahi Techno Glass Co., Shizuoka, Japan) was performed via measurement of the stimulated emission of orange light (500–700 nm) from the dosimeter when a 365-nm mono-energetic laser was exposed on the irradiated dosimeter. During dose readout, there were two readout modes, based on dose values: low-dose-range mode (10 μGy–10 Gy), and high-dose-range mode (1–500 Gy). In this study, we used the high-dose-range mode. The advantages of the RPLGD, in comparison with the TLD, include its good reproducibility (1%) and relatively low energy dependency at energies higher than 200 keV (23-25). Additionally, in comparison with the TLD or OSLD, the RPLGD has a smaller incident-beam-angle dependency and lower toxicity inside the human body (26-29). The RPLGD is comparable to the Al_2_O_3_:C OSLD, which is made of a luminescent material similar to that of TLD, but has a different excitation source and uses a different readout technique. The OSLD is not necessary for the heating procedure, but its detector is affected by visible light. A variety of studies have attempted to characterize the dose response of OSLD; the supra-linearity of the dose response has been reported to be above 300 cGy, with angular dependency and relatively low energy dependency, when a clinical mega-voltage photon and electron beam were used (29). In 2010, Jursinic reported that the OSLD response to dose was supralinear when the detector received accumulative doses, and that a dose response accuracy of ±0.5% could be achieved if the sensitivity and extent of supra-linearity were established for each OSLD (30).

### Experimental measurements

Figure 1 illustrates the experimental setup. Forty-five RPLGDs were irradiated on a 10 × 10 cm^2^ open field, using 6, 10, and 15 MV photon beams. The source-to-surface distance (SSD) was 100 cm, and the depths of the detectors were set relative to the dose-maximum depth of energy: 1.5 cm (6 MV), 2.5 cm (10 MV), and 3.0 cm (15 MV). A 20-cm-thick block of water-equivalent solid phantom was placed behind the detectors to gather the backscatter of radiation. A 1-cm-thick homemade RPLGD phantom was inserted between the build-up phantom and backscatter phantom. Although the phantom was designed to contain up to 42 RPLGDs, in this study, only 3 RPLGDs were inserted at a time into the homemade phantom to be irradiated with radiation. For accurate dose delivery, the monitor unit (MU) value was corrected, considering the output factor of the linear accelerator measured according to TRS-398 (31) and the percentage depth dose (PDD) of the beam data.

**Figure 1.**
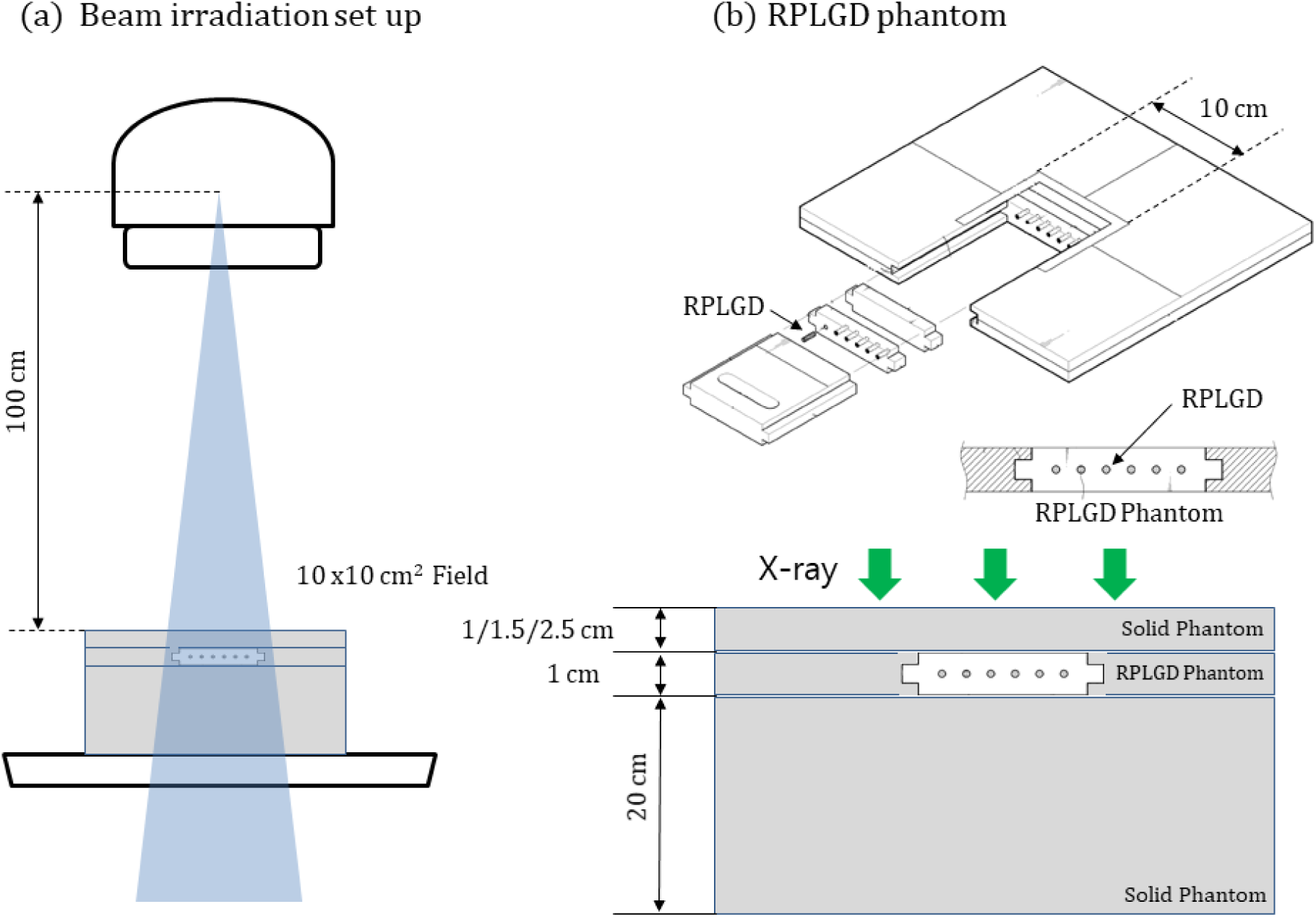
Measurement setup. **(a) Beam irradiation setup (SSD: 100 cm, field size: 10 × 10 cm**^**2**^, **Depth: Dmax), (b) Homemade RPLGD phantom**.

For each energy level, five groups were created, wherein each group involved three detectors and its own set of experimental conditions. For most of the groups, except Group 2, the detectors were initialized each time by being annealed at 400 °C for 1 h before each irradiation. For Group 2, the annealing process was skipped during the measurements to evaluate the effect of the annealing process.

Groups 1 and 2 were irradiated to values of 1 Gy, 5 Gy, 10 Gy, 50 Gy, and then 100 Gy, in order; readings were taken after each irradiation to estimate the dose readout for the integral dose, with (Group 1) and without (Group 2) the annealing procedure. For Group 3, the RPLGD reading procedure was performed after irradiation with a dose of approximately 10 Gy. This procedure was repeated 10 times to evaluate the fading effect of RPLGD. For Groups 4 and 5, the RPLGD reading procedure was conducted after irradiation with 50 Gy (Group 4) and 100 Gy (Group 5). In the same way as with Group 3, these procedures were repeated 10 times for each group at three different levels of energy: 6, 10, and 15 MV.

## Results

Figure 2 depicts the measurement results for Groups 1 (dashed line) and 2 (solid line). For each run with Group 1, the setup was annealed at 400 °C for 1 h before each irradiation. The annealing process was skipped for Group 2. Each group was irradiated to 1, 5, 10, 50, and 100 Gy, in order. The energies of the irradiated beam were (a) 6 MV (open circles: annealed; shaded circles: accumulated), (b) 10 MV (open triangles: annealed; shaded triangles: accumulated), and (c) 15 MV (open squares: annealed; shaded squares: accumulated). Figure 2(d) depicts the dose ratios of measurements and irradiation doses. The solid red lines in Figures 2(a), 2(b), and 2(c) represent the expected values when the reading of RPLGD increases in proportion to the irradiation dose. Group 2 exhibits a hyperlinear response, as depicted in Figure 2. In Group 1, when the doses were sequentially increased to 1, 5, 10, 50, and 100 Gy over a total of 5 times, the dose response (dashed black line) did not exhibit a significant difference in comparison with the expected value (solid red line); however, in Group 2, the dose response (solid black line) was higher overall. Table 1 presents the percentages of the measured values relative to the irradiation doses for Groups 1 and 2. The top three rows list the results for Group 1, wherein the setup was initialized each time by being annealed at 400 °C for 1 h before each irradiation. Meanwhile, the bottom three rows list the results for Group 2, wherein the setup was not annealed. In the case of Group 1, the percentage of the measured value relative to the beam irradiation was estimated to be within 5%, whereas for Group 2, the percentages of the measured values relative to the beam irradiation were 118.7 ± 1.9% for 6 MV, 112.2 ± 2.7% for 10 MV, and 101.5 ± 2.3% for 15 MV.

**Table 1.** Measurement results for Groups 1 and 2. The top three rows list the results for Group 1, wherein the setup is initialized each time by being annealed at 400 °C for 1 h before each irradiation. The bottom three rows list the results for Group 2, wherein the setup is not annealed. The irradiation dose is sequentially increased to 1, 5, 10, 50, and 100 Gy, over a total of five steps

| Annealing | Energy<br>(MV) | Exposed Dose (Gy) |  |  |  |  |
| --- | --- | --- | --- | --- | --- | --- |
|  |  | 1 | 5 | 10 | 50 | 100 |
| **Yes**<br>**(Group 1)** | 6 | $100.0 \pm 0.5$ | $98.6 \pm 0.4$ | $101.4 \pm 0.3$ | $96.2 \pm 0.3$ | $104.0 \pm 0.3$ |
| | 10 | $100.0 \pm 1.6$ | $99.9 \pm 0.9$ | $102.4 \pm 1.4$ | $97.3 \pm 0.9$ | $104.6 \pm 1.9$ |
| | 15 | $100.0 \pm 1.0$ | $102.7 \pm 1.9$ | $97.0 \pm 1.5$ | $92.9 \pm 2.3$ | $95.0 \pm 2.6$ |
| **No**<br>**(Group 2)** | 6 | $100.0 \pm 0.3$ | $99.0 \pm 0.9$ | $102.4 \pm 0.8$ | $98.7 \pm 3.1$ | $118.7 \pm 1.9$ |
| | 10 | $100.0 \pm 0.9$ | $94.8 \pm 0.8$ | $98.9 \pm 0.8$ | $98.1 \pm 0.2$ | $112.2 \pm 2.7$ |
| | 15 | $100.0 \pm 0.7$ | $102.3 \pm 1.0$ | $90.1 \pm 2.4$ | $93.3 \pm 1.4$ | $101.5 \pm 2.3$ |

**Figure 2.**
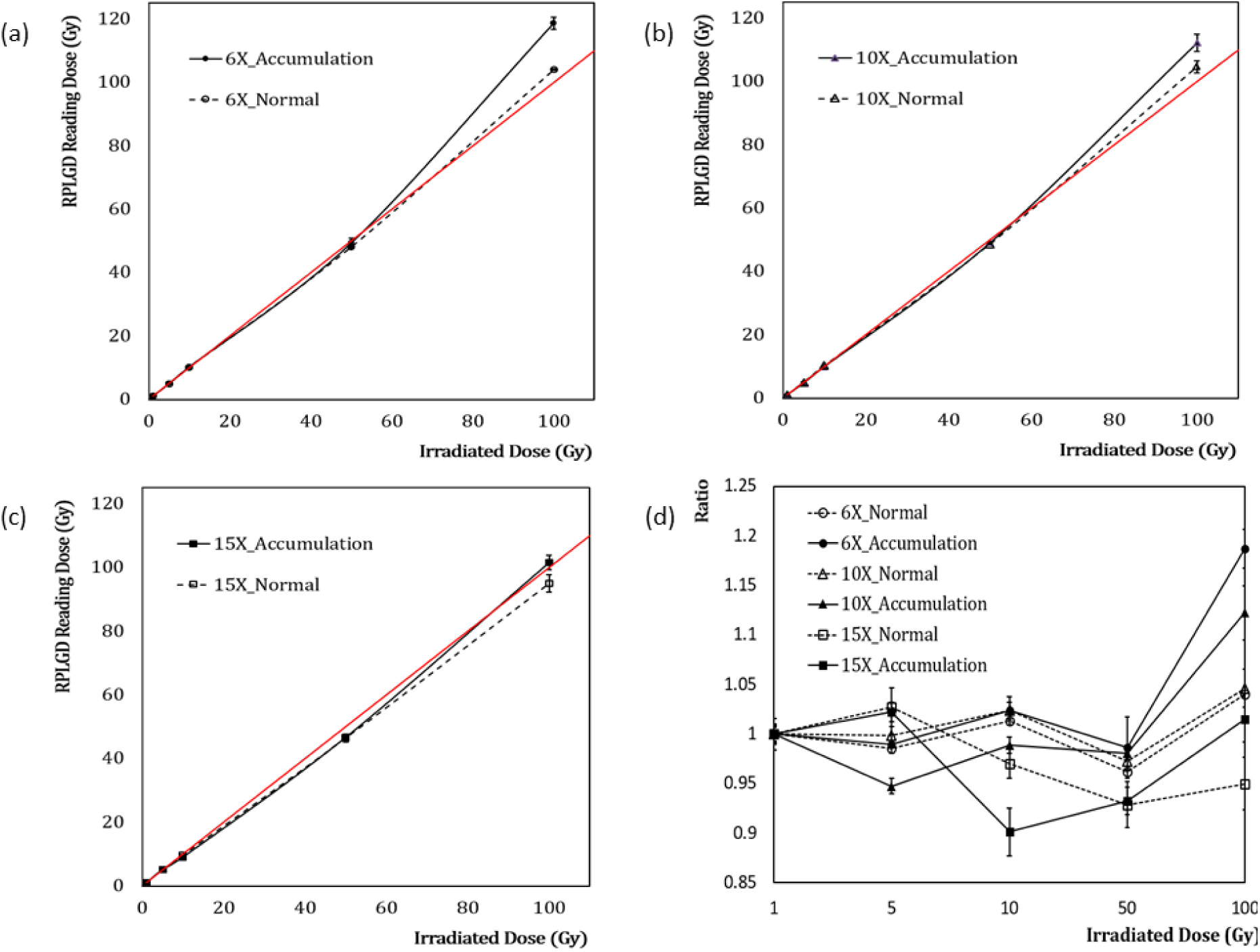
Measurement results for Group 1 (dashed line) and Group 2 (solid line). For each run with Group 1, the setup is annealed at 400 °C for 1 h before each irradiation. The annealing process is skipped for Group 2. Each group is irradiated to 1, 5, 10, 50, and 100 Gy, in order. The energies of the irradiated beam are (a) 6 MV (open circles: annealed; shaded circles: accumulated), (b) 10 MV (open triangles: annealed; shaded triangles: accumulated), and (c) 15 MV (open squares: annealed; shaded squares: accumulated). (d) Dose ratios of measurements and irradiation doses

Figure 3 depicts the measurement results for (a) Groups 3, (b) 4, and (c) 5, assuming that the sensors were reused for similar doses over time. In the experiment, wherein the sequence of initialization, beam irradiation, and reading was repeated 10 times, the responsiveness of the RPLGD gradually decreased as the number of repetitions increased. Table 2 presents the measurement results with respect to the number of irradiations, from 1 to 10 times. Each group was irradiated at doses of 10, 50, and 100 Gy, with three different energies, namely, 6, 10, and 15 MV. The values are normalized percentages with respect to the 1st measurement values. The measurement results after 10 repeated beam irradiations were 84.6 ± 1.9%, 87.5 ± 2.4%, and 93.0 ± 3.0% at 6 MV, 10 MV, and 15 MV, respectively.

**Table 2.** Measurement results for 10 irradiations for evaluating the fading effect of RPLGD. Sensors are assumed to have been reused for similar doses over time. Each group is irradiated at doses of 10, 50, and 100 Gy, with three different energies, namely, 6, 10, and 15 MV. The values are normalized percentages with respect to the 1^st^ measurement values.

| Energy<br>(MV) | Dose<br>(Gy) | Sequence |  |  |  |  |  |  |  |  |  |
| --- | --- | --- | --- | --- | --- | --- | --- | --- | --- | --- | --- |
|  |  | 1 | 2 | 3 | 4 | 5 | 6 | 7 | 8 | 9 | 10 |
| 6 | 10 | 100.0 | 97.3 | 97.6 | 96.5 | 102.0 | 96.0 | 90.4 | 87.6 | 86.1 | 85.9 |
| | | $\pm 0.0$ | $\pm 1.2$ | $\pm 2.0$ | $\pm 0.9$ | $\pm 1.8$ | $\pm 2.4$ | $\pm 1.4$ | $\pm 1.9$ | $\pm 1.0$ | $\pm 2.1$ |
|  | 50 | 100.0 | 96.6 | 96.9 | 94.9 | 98.2 | 95.1 | 92.6 | 85.8 | 85.7 | 83.5 |
| | | $\pm 0.0$ | $\pm 0.5$ | $\pm 0.7$ | $\pm 0.4$ | $\pm 0.5$ | $\pm 0.6$ | $\pm 1.7$ | $\pm 0.8$ | $\pm 2.0$ | $\pm 1.0$ |
|  | 100 | 100.0 | 94.6 | 95.8 | 94.9 | 97.2 | 92.3 | 88.1 | 85.7 | 82.6 | 84.4 |
| | | $\pm 0.0$ | $\pm 1.6$ | $\pm 1.5$ | $\pm 0.8$ | $\pm 1.9$ | $\pm 2.4$ | $\pm 1.7$ | $\pm 2.8$ | $\pm 2.1$ | $\pm 1.8$ |
| 10 | 10 | 100.0 | 100.4 | 98.8 | 98.4 | 99.4 | 91.3 | 98.1 | 90.1 | 88.5 | 88.2 |
| | | $\pm 0.0$ | $\pm 1.0$ | $\pm 0.6$ | $\pm 0.7$ | $\pm 2.2$ | $\pm 0.5$ | $\pm 2.0$ | $\pm 1.9$ | $\pm 2.5$ | $\pm 2.4$ |
|  | 50 | 100.0 | 100.3 | 98.9 | 98.4 | 100.1 | 94.7 | 99.0 | 90.3 | 89.2 | 89.0 |
| | | $\pm 0.0$ | $\pm 1.5$ | $\pm 0.7$ | $\pm 0.9$ | $\pm 0.5$ | $\pm 1.0$ | $\pm 0.9$ | $\pm 0.3$ | $\pm 0.5$ | $\pm 1.1$ |
|  | 100 | 100.0 | 97.6 | 96.0 | 95.7 | 97.7 | 95.7 | 90.7 | 88.3 | 85.5 | 85.4 |
| | | $\pm 0.0$ | $\pm 1.7$ | $\pm 2.5$ | $\pm 0.3$ | $\pm 0.9$ | $\pm 0.9$ | $\pm 0.7$ | $\pm 1.1$ | $\pm 0.1$ | $\pm 1.7$ |
| 15 | 10 | 100.0 | 100.1 | 96.9 | 91.4 | 90.3 | 93.1 | 98.2 | 93.4 | 92.8 | 92.1 |
| | | $\pm 0.0$ | $\pm 1.8$ | $\pm 4.0$ | $\pm 4.0$ | $\pm 1.5$ | $\pm 2.9$ | $\pm 2.8$ | $\pm 3.4$ | $\pm 2.7$ | $\pm 1.8$ |
|  | 50 | 100.0 | 99.7 | 98.1 | 95.5 | 95.2 | 95.5 | 97.0 | 96.7 | 94.2 | 95.1 |
| | | $\pm 0.0$ | $\pm 0.7$ | $\pm 0.9$ | $\pm 0.4$ | $\pm 0.5$ | $\pm 1.3$ | $\pm 1.4$ | $\pm 0.5$ | $\pm 2.1$ | $\pm 1.7$ |
|  | 100 | 100.0 | 98.4 | 98.5 | 94.4 | 94.9 | 94.5 | 92.8 | 96.6 | 93.1 | 92.0 |
| | | $\pm 0.0$ | $\pm 1.6$ | $\pm 1.1$ | $\pm 1.5$ | $\pm 0.7$ | $\pm 1.6$ | $\pm 1.7$ | $\pm 2.1$ | $\pm 2.0$ | $\pm 3.9$ |

**Figure 3.**
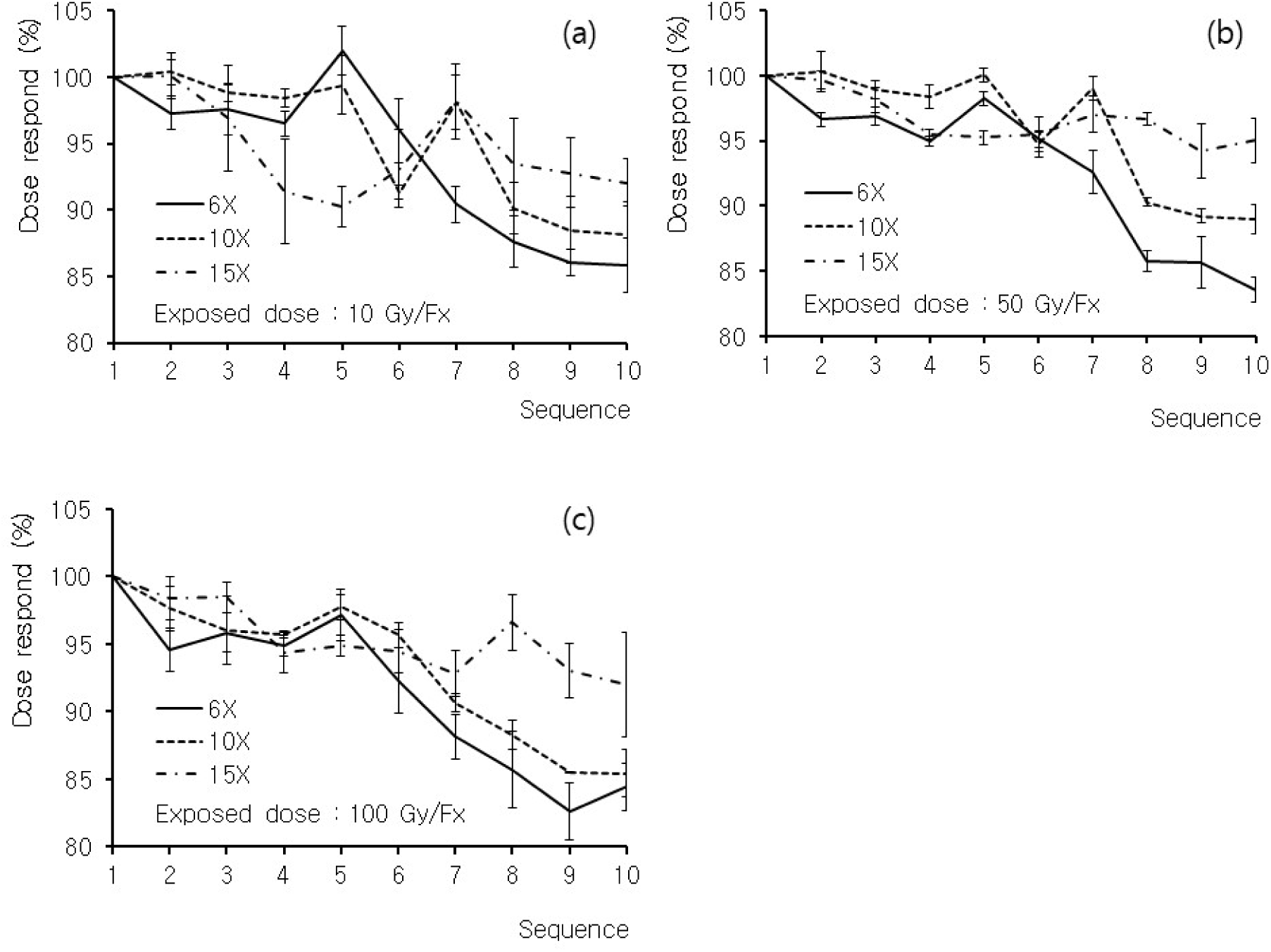
Measurement results for Groups 3, 4, and 5. Sensors are assumed to have been reused for similar doses over time. Groups 3, 4, and 5 are irradiated at doses of (a) 10 Gy, (b) 50 Gy, and (c) 100 Gy, respectively. Each group is irradiated with three different energies: 6 MV (solid line), 10 MV (dashed line), and 15 MV (dash-dotted line).

## Discussion

Figure 2 depicts that, up to a 50 Gy irradiation, the dose response was not significantly different; however, after a 100 Gy irradiation, the response in Group 2 increased to 118%. Thus, when measured with the RPLGD, the dosimeter response was not proportional to the irradiation dose and exhibited a hyperlinear pattern that resulted in a slightly higher reading when the setup was cumulatively irradiated without annealing. Additionally, the hyperlinear response of Group 2 did not appear significantly until the setup was irradiated with 50 Gy, but exhibited a significant increase in dose response by up to 118% when the setup was irradiated with 100 Gy. As presented in Table 1, in the case of Group 1, the percentage of the measured value relative to beam irradiation was estimated to be within 5%, whereas for Group 2, the percentages of the measured values relative to beam irradiation were 118.7 ± 1.9% for 6 MV, 112.2 ± 2.7% for 10 MV, and 101.5 ± 2.3% for 15 MV.

Furthermore, the changing effect in dose response due to dose accumulation was also dependent on beam energy. As shown in Figure 2 and Table 1, the data for the 15 MV beam dose do not exhibit hyperlinearity in both the annealed (Group 1) and accumulated (Group 2) cases. Table 3 presents the fitting results for the annealed (Group 1) and accumulated (Group 2) cases via the least square function and quadratic polynomial function. Data fitting was performed using SigmaPlot (Sigmaplot version 14.0, Systate Software, Inc. San Jose, CA 95131, USA) with the Shapiro–Wilk fitting evaluation tool. For the annealed (Group 1) case, the P values obtained from the Shapiro–Wilk test for most of the data were greater than 0.05 with the least square function. For the non-annealed (Group 2) case, the P values for the 6 MV and 10 MV data were lesser than 0.05 with the least square function. For the 15 MV non-annealed data (Group 2), the P value from the Shapiro–Wilk test was 0.6429.

**Table 3.**
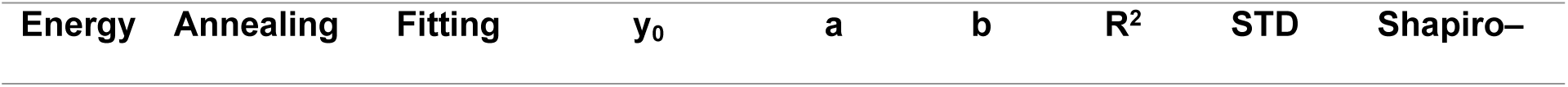
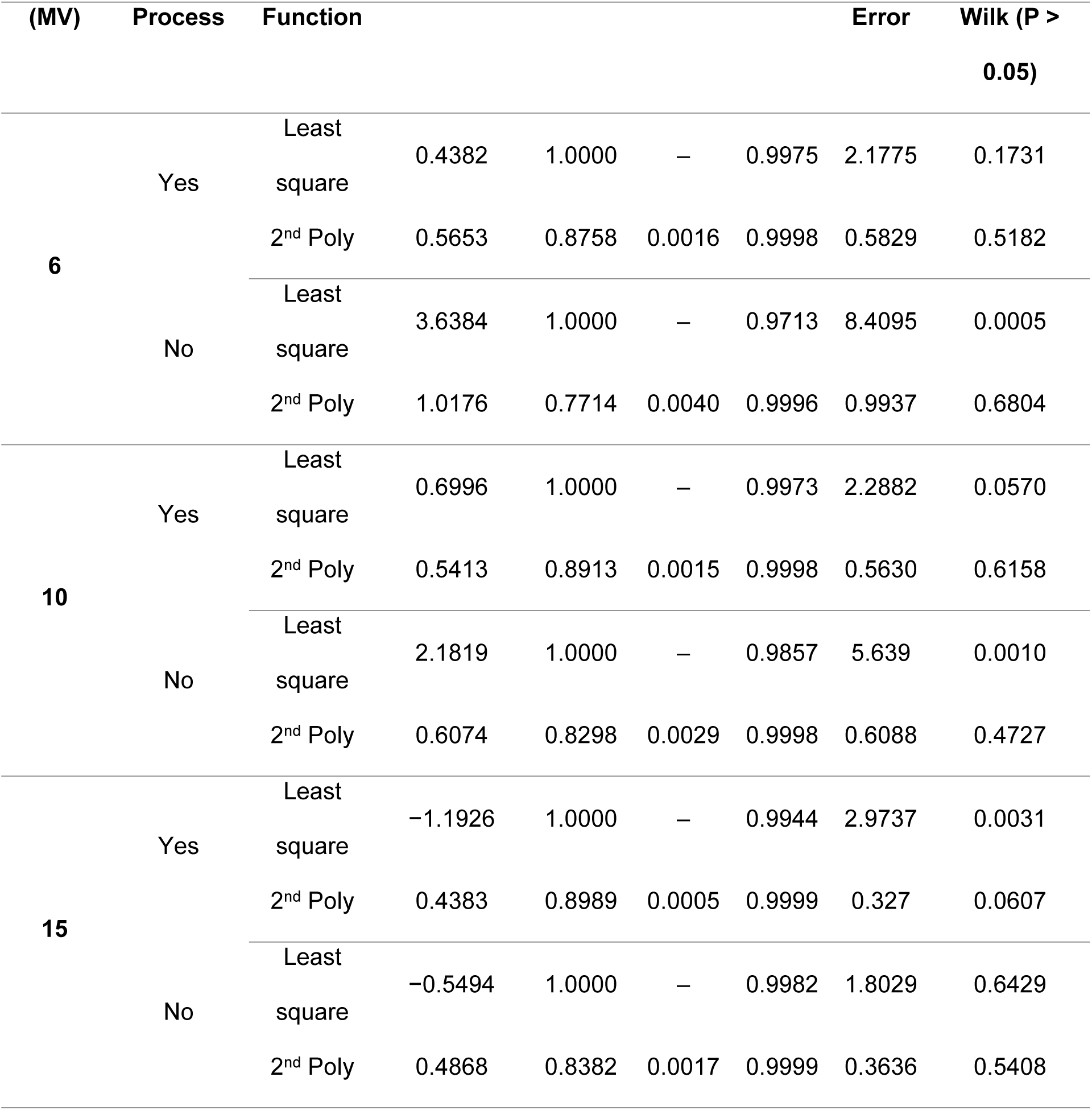
Fitting results for Groups 1 and 2. Group 1 is initialized each time via annealing at 400 °C for 1 h before each irradiation, whereas Group 2 is not annealed. The irradiation dose is sequentially increased to 1, 5, 10, 50, and 100 Gy, over a total of five steps. The least squares function and quadratic polynomial function are used as fitting functions for the measurement results.

The fitting results also demonstrated that the dose response for the annealed case (Group 1) was linear with respect to the irradiation dose. For the non-annealed case (Group 2), the 6 MV and 10 MV data exhibited a hyperlinear dose response, whereas the 15 MV data exhibited a linear response. Therefore, when measurements are being conducted using RPLGD, particularly in the case of measurements at low energies with high doses, it is suggested that the annealing process be performed each time to reduce errors due to dose accumulation. In contrast, it is possible to correct the data response by considering the hyperlinearity of RPLGD through the use of a fitting parameter. In the results for Groups 1 and 2, the dispersion between the measuring elements was measured to be within 3%. Therefore, given that, the difference between the sensors is small in the measurement, it is possible to correct the read dose value via acquisition of factors related to the cumulative dose response of each device with respect to energy, rather than via initialization of the sensor every time before measurement.

Figure 3 depicts that the responsiveness of the RPLGD gradually decreases to approximately –15% as the number of repetitions increases. This fading phenomenon is more prominent at lower energies than at higher energies. However, a significant relationship does not appear to exist between the magnitude of the dose irradiated to the device and the fading effect. As presented in Table 4, the fading slope obtained via a least square fit was largest at 6 MV and smallest at 15 MV. The measurement results after 10 repeated beam irradiations were 84.6 ± 1.9%, 87.5 ± 2.4%, and 93.0 ± 3.0%, at 6 MV, 10 MV, and 15 MV, respectively. However, until the fifth measurement in the overall measurement process, the fading effect was insignificant, being within 5%. Therefore, if possible, recalibrating the RPLGD after five uses is necessary to correct the sensitivity degradation due to the fading effect. Although correcting the measured value by considering the effect of fading on each RPLGD is possible at each energy level, in this case, the similarity of the dose responses among the RPLGDs should be evaluated.

**Table 4.**
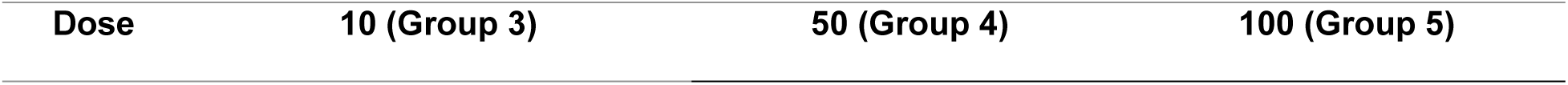
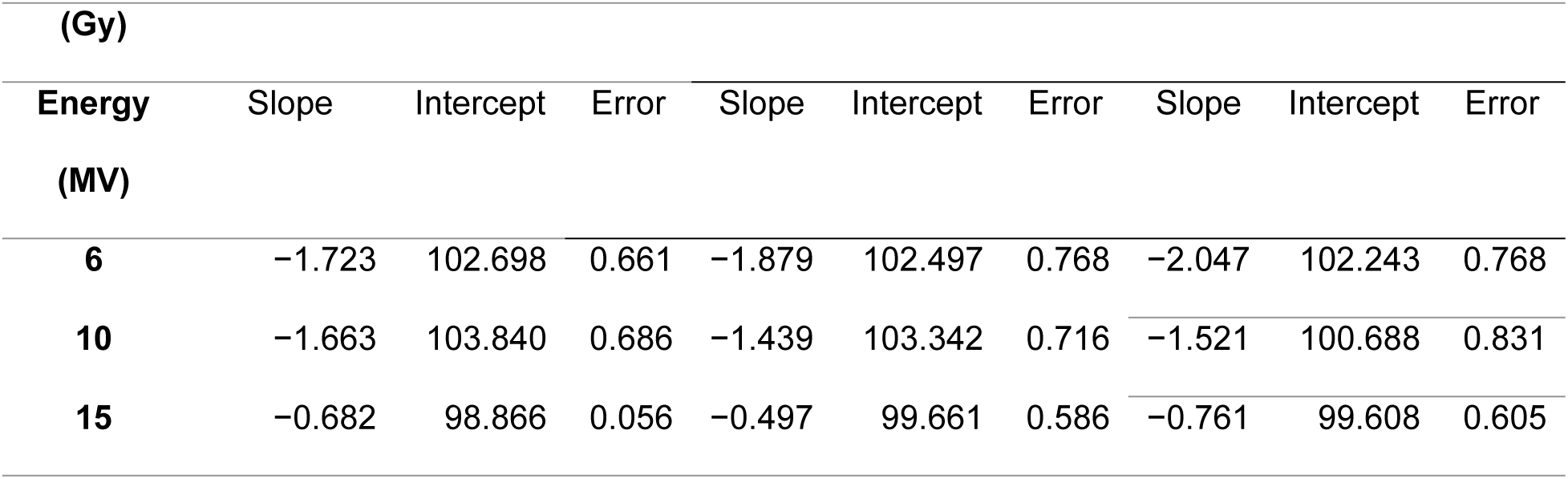
Fitting results for fading slope, intercept, and error obtained via least square fit. These results are for three values of energy and three values of dose. Groups 3, 4, and 5 are irradiated with doses of 10, 50, and 100 Gy, respectively. Each group is irradiated at three different energies (6, 10, and 15 MV).

| Dose | 10 (Group 3) | 50 (Group 4) | 100 (Group 5) |
| --- | --- | --- | --- |

| (Gy) |  |  |  |  |  |  |  |  |  |
| --- | --- | --- | --- | --- | --- | --- | --- | --- | --- |
| Energy | Slope | Intercept | Error | Slope | Intercept | Error | Slope | Intercept | Error |
| (MV) |  |  |  |  |  |  |  |  |  |
| 6 | -1.723 | 102.698 | 0.661 | -1.879 | 102.497 | 0.768 | -2.047 | 102.243 | 0.768 |
| 10 | -1.663 | 103.840 | 0.686 | -1.439 | 103.342 | 0.716 | -1.521 | 100.688 | 0.831 |
| 15 | -0.682 | 98.866 | 0.056 | -0.497 | 99.661 | 0.586 | -0.761 | 99.608 | 0.605 |

In this study, measurements were performed using only three devices in each group; however, more detailed results can be obtained through the use of a greater number of glass dosimeters to reduce the statistical uncertainty in the near future.

## Conclusions

The non-annealed RPLGD response to dose was determined to be hyperlinear for the 6 MV and 10 MV photon beams but not for the 15 MV photon beam. This response was also found to change according to the amount of accumulated dose delivered to the RPLGD. Additionally, the annealed RPLGD was observed to exhibit a fading phenomenon when the measurement was repeated several times, and the fading effect was relatively significant at low energies in comparison with high energies. In this study, measurements were performed using only three devices in each group; however, more detailed results and predictions for the responses can be obtained in the near future through the use of a greater number of glass dosimeters to reduce statistical uncertainty. Moreover, after up to five repeated uses, the calibration of each device needs to be performed to reduce the uncertainty caused by the fading effect. Furthermore, we suggest that the annealing procedure should be performed before each measurement.

## Acknowledgments

This work was supported by the general researcher program (NRF-2018R1D1A1B07050217) and the nuclear safety research program (No. 2003013-0120-CG100) through the Korea Foundation of Nuclear Safety (KOFONS), using the financial resource granted by the Nuclear Safety and Security Commission (NSSC), Republic of Korea

